# Computational identification of key transcription factors for embryonic and postnatal *Sox2+* dental epithelial stem cell

**DOI:** 10.1101/2023.12.22.573158

**Authors:** Fan Shao, Eric Van Otterloo, Huojun Cao

**Affiliations:** Iowa Institute for Oral Health Research, University of Iowa College of Dentistry and Dental Clinics, Iowa City, IA 52242; Division of Biostatistics and Computational Biology, University of Iowa College of Dentistry and Dental Clinics, Iowa City, IA 52242; Department of Endodontics, University of Iowa College of Dentistry and Dental Clinics, Iowa City, IA 52242; Department of Periodontics, University of Iowa College of Dentistry and Dental Clinics, Iowa City, IA 52242; Department of Anatomy and Cell Biology, University of Iowa Carver College of Medicine, Iowa City, IA 52242

**Author notes:** Correspondence should be addressed to H. C. Division of Biostatistics and Computational Biology Department of Endodontics Iowa Institute for Oral Health Research University of Iowa College of Dentistry, Iowa City, 52242.

## Abstract

While many reptiles can replace their tooth throughout life, human loss the tooth replacement capability after formation of the permanent teeth. It was thought that the difference in tooth regeneration capability depends on the persistence of a specialized dental epithelial structure, the dental lamina that contains dental epithelial stem cells (DESC). Currently, we know very little about DESC such as what genes are expressed or its chromatin accessibility profile. Multiple markers of DESC have been proposed such as *Sox2* and *Lgr5*. Few single cell RNA-seq experiments have been performed previously, but no obvious DESC cluster was identified in these scRNA-seq datasets, possible due to that the expression level of DESC markers such as *Sox2* and *Lgr5* is too low or the percentage of DESC is too low in whole tooth. We utilize a mouse line *Sox2-GFP* to enrich *Sox2+* DESC and use Smart-Seq2 protocol and ATAC-seq protocol to generate transcriptome profile and chromatin accessibility profile of P2 *Sox2+* DESC. Additionally, we generate transcriptome profile and chromatin accessibility profile of E11.5 Sox2+ dental lamina cells. With transcriptome profile and chromatin accessibility profile, we systematically identify potential key transcription factors for E11.5 *Sox2+* cells and P2 *Sox2+* cells. We identified transcription factors including *Pitx2, Id3, Pitx1, Tbx1, Trp63, Nkx2-3, Grhl3, Dlx2, Runx1, Nfix, Zfp536*, etc potentially formed the core transcriptional regulatory networks of *Sox2+* DESC in both embryonic and postnatal stages.

## INTRODUCTION

Many reptiles, reiteratively replace their teeth throughout life (polyphyodonts)[1–10]. In contrast, human are characterized by replacement of teeth once throughout life (diphyodonts). The development of teeth is governed by reciprocal interactions between epithelium and the underlying neural crest derived mesenchyme. Multiple postnatal human dental mesenchymal stem cells have been isolated such as dental pulp stem cells (DPSCs), stem cells from exfoliated deciduous teeth (SHED), and dental follicle progenitor cells (DFPCs)[11–14]. In contrast, there is no available source of human dental epithelial stem cells (DESC) after permanent teeth were formed. It has been thought that the difference in tooth regeneration capability between different species depends on the persistence of a specialized epithelial structure, the dental lamina (DL), which contains dental epithelial stem cells[1]. The DL of diphyodonts (e.g., human) degrades after initiation of permanent teeth while the DL of polyphyodonts (e.g., shark, reptiles) persists throughout life. It has been shown that prolonged survival of the DL in humans leads to supernumerary tooth formation[15, 16].

It has been shown that in most species, including mice and humans, tooth development starts with a localized thickening of the maxillary and mandibular ectoderm, resulting in a ‘strip’ of dental epithelium (i.e., the DL) that contains DESC and have the capability to induce a tooth development program[17]. All teeth will originate within this DL ‘strip’, going through a series of developmental stages including: placode, bud, cap, bell, etc. In mice, DL strip formed at embryonic day 11.5 (E11.5). We have previously found that SOX2 and TFAP2B expression broadly divided the mandibular epithelium into two complementary domains along the dorsal-ventral axis. Specifically, at E11.5, SOX2 expression covering posterior domain epithelium and most cells in DL, while TFAP2B expression being high throughout the ventral side of the mandibular epithelium, was dramatically reduced after reaching SOX2 positive cells in the DL[18]. In mouse incisor, we can find SOX2 expression become gradually restricted and finally only expressed at selected cells in labial cervical loop region (dental stem cell niche). The mouse incisor presents as a power model for studying DESC as it contains DESC that enable continuously growth of incisor.

Multiple markers of DESC have been proposed including: *Sox2, Lgr5, Gli1, Lrig1, Bmi1* and *Ptch1* [19–24]. Among them, *Sox2* is the most well characterized marker of DESC. It has been shown that *Sox2* positive cell give rise to all epithelial cell lineages in tooth[19, 25]. Additionally, so far, *Sox2* seems expressed in DL, which contains DESC, of all species that has been examined (including polyphyodonts and diphyodonts)[9, 19, 25–28]. It is possible that there are multiple populations of dental epithelial stem cell that can give rise multiple/all epithelial cell lineages in the DL of polyphyodonts or labial cervical loop of mouse incisor, just like other tissues’ epithelial stem cell niches such as hair follicle or small intestine. However, currently, we still know very little about DESC. Several single cell RNA-seq experiments has been performed with mouse and human tooth. However, so far, it seems that scRNA-seq fail to capture the distinct DESC cluster that were marked by the abovementioned known DESC markers such as *Sox2, Lgr5*, etc[24, 29–31]. This failure could be caused by: 1) the occurrence of ‘dropouts’ problem for genes that are not highly expressed in current scRNA-seq technologies; 2) the frequency of DESC is too low to be able to get enough cell number in previous experiments (without enrichment with DESC markers). To address these two issues, in this study, we utilize a mouse line *Sox2-GFP*[32] in which GFP expression is driven by endogenous promoter of *Sox2*, to enrich DESC; additionally we use Smart-Seq2 protocol, which has been shown have much improved efficiency in profiling medium/low expressed genes comparing with other scRNA-seq technologies[33]. Besides transcriptome profiles, we also generated chromatin accessibility profiles of *Sox2+* DESC cells with ATAC-seq assay[34]. We compared the transcriptome and chromatin accessibility profiles of *Sox2+* DESC cells in two stages (E11.5 and P2) and identified potential key transcription factors that are shared or different in these two stages.

## RESULTS

To generate transcriptome of *Sox2+* dental epithelial stem cell, we utilize a mouse line *Sox2-GFP*[32] in which GFP expression is driven by endogenous promoter of *Sox2*. Labial cervical loop (LaCL) region of Postnatal day 2 (P2) lower incisors were dissected and dissociated into single cells. Based on GFP expression, we collected *Sox2* positive cells (*Sox2+*; enriched for dental epithelial stem cells) and *Sox2* negative cells (*Sox2-*; contain mixture of cell populations such as differentiating epithelial cell and mesenchymal cells) (**Fig. 1A**). To identify genes that are enriched or upregulated in *Sox2+* dental epithelial stem cell, we generated transcriptome profiles of *Sox2+* cells and *Sox2-* cells with Smart-Seq2 RNA-seq protocol[33].

**Figure 1.**
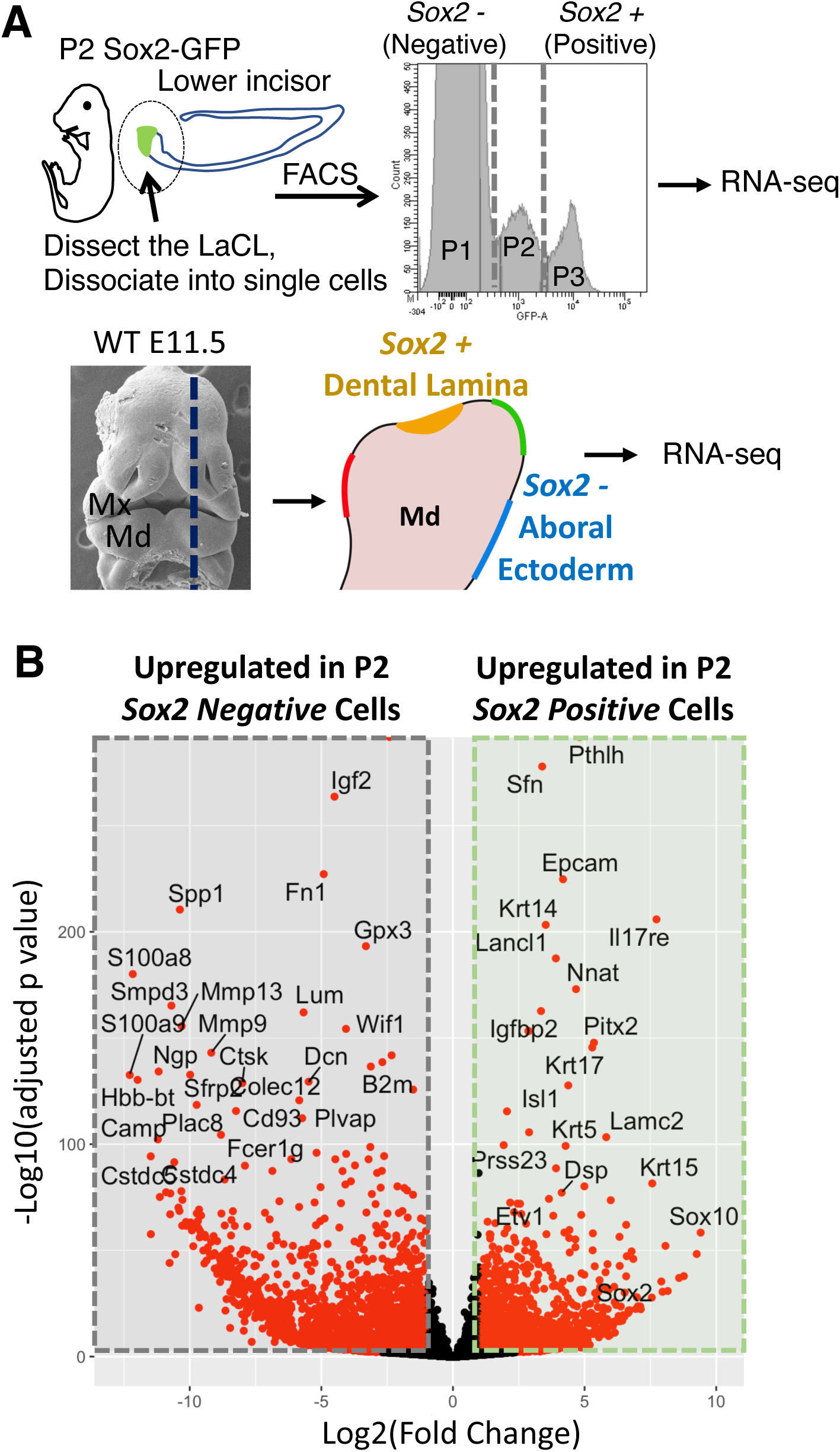
Transcriptome profiling of the P2 *Sox2+* dental epithelial stem cell and E11.5 *Sox2+* dental lamina cells. **A)** Workflow of tissue preparation for RNA-seq experiment. For P2 samples, *Sox2+* cells and Sox2-cells were isolated from lower incisor labial cervical loop. For E11.5 samples, Laser-Microdissection Microscopy were used to isolated *Sox2+* dental lamina cells and *Sox2-* aboral ectoderm cells. **B)** Volcano plot showing differentially expressed genes (DEGs) of the P2 *Sox2+* cells compared with P2 *Sox2-* cells. Red dots indicate DEGs with a absolute log2 fold-change larger than 1.0 and a adjusted p value less than 0.01. Abbreviations: DL, dental lamina; AE, aboral ectoderm; Md, Mandible; Mx, Maxilla.

We performed differentially expressed gene (DEG) analysis between the transcriptome profiles of these two populations with DESeq2[35]. To control for variability associated with lowly expressed genes, we further applied the ASHR algorithm[36] to estimate shrunken (conservative) log2 fold changes. Through this analysis, we identified 2072 genes were significantly (adjusted P value < 0.01 and |log2 fold changes| > 1.0) upregulated in *Sox2+* cells comparing with *Sox2-* cells in P2 lower incisor labial cervical loop (**Fig. 1B**). Similarly, we identified 3040 genes were significantly upregulated in Sox2-cells (comparing with *Sox2+* cells). In *Sox2-* cells, we found upregulation of genes critical for ameloblast cell differentiation such as *Mmp13* and *Mmp9* (**Fig. 1B**). Additionally, because *Sox2-* cells contain mixture of cell populations that includes mesenchymal cells, we found epithelial and mesenchymal cell markers among the top differentially expressed gene (for example: *Epcam, Fn1, Krt14, Krt17, Krt5* and *Krt15*; **Fig. 1B**).

In DEG analysis comparing *Sox2+* cell with *Sox2-* cell from P2 lower incisor labial cervical loop, we found many top upregulated (based on P value and log2 fold change) genes in *Sox2+* cells are transcription factors and signal pathways genes. As expected, *Sox2* is one of the top upregulated genes (**Fig. 1B**). Besides *Sox2*, we found transcription factors including: *Pitx2, Isl1, Etv1* and *Sox10* among top upregulated genes (**Fig. 1B**). By intersected the DEG results with all known mouse TFs listed in the AnimalTFDB database[37], we identified total 105 transcription factors that are significantly upregulated in *Sox2+* cells comparing with *Sox2*-cells. Besides transcription factors, we also few regulators of signal pathways among the top DEGs. Specifically, we found *Pthlh* and Igfbp2 are significantly upregulated in *Sox2+* cells, while *Igf2* and *Sfrp2* are significantly downregulated in *Sox2+* cells.

In most species, including mice and humans, tooth development starts with a localized thickening of the maxillary and mandibular ectoderm, resulting in a ‘strip’ of dental epithelium (i.e., the DL) that have the capability to induce a tooth development program[17]. In mice, DL formed at Embryonic day 11.5 (E11.5). We have previously compared transcriptome profiles of four domains along the dorsal-ventral axis of the E11.5 mouse mandibular epithelium[18]. Specifically, in pair-wise DEG analysis comparing E11.5 dental lamina (*Sox2+* DL cell, which give rise all dental epithelial cells) with E11.5 aboral ectoderm (*Sox2-* AE cell, which give rise facial skin epithelial cells), we found 117 genes significantly upregulated and 242 genes significantly downregulated in DL. To determine the overlapping between DEG results from these two stages (E11.5 and P2) comparing *Sox2+* cells with *Sox2-* cells, we first intergrated log2 fold change in these two comparision (**Fig. 2A**). We found few transcription factors were consistenly among the top upregulated genes in both stages including: *Sox2, Pitx2, Nkx2-3, Etv5* and *Pitx1*(**Fig. 2A**). On the contrary, we found that few transcription factors that was upregulated in only one stage. For example, *Six1* and *Zfp536* were significant upregulated in E11.5 *Sox2+* cells (DL) when comparing with E11.5 *Sox2-* cells, but at P2 stage, their expression is not significantly different when comparing with *Sox2+* cells with *Sox2-* cells. Similarly, we found *Sox10*’s expression become significantly enriched in P2 stage *Sox2+* cells while no significant difference was found at E11.5 stage between *Sox2+* cells and *Sox2-* cells (**Fig. 2A**).

**Figure 2.**
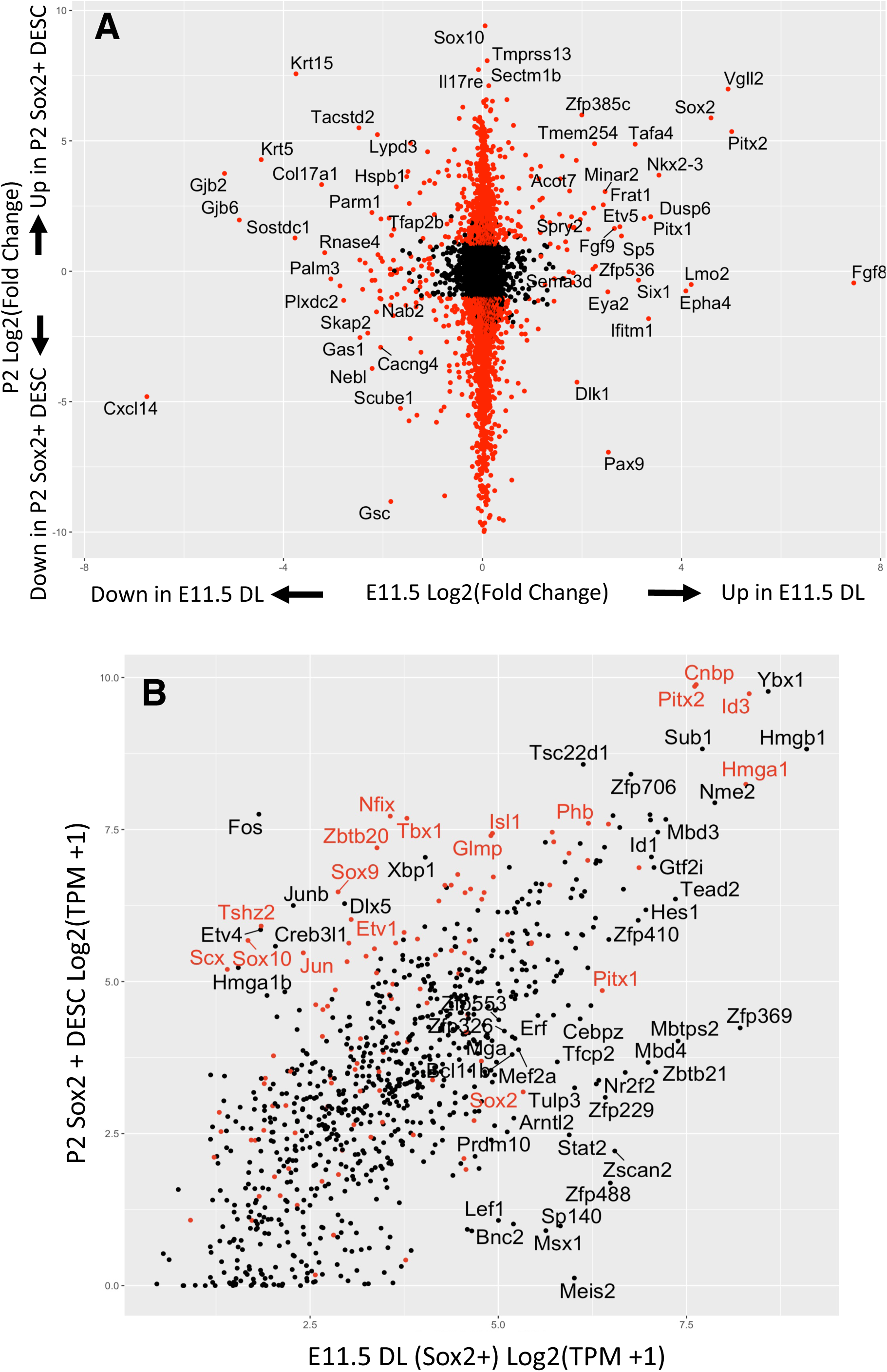
Integration and comparison of P2 *Sox2+* DESC and E11.5 *Sox2+* DL cells. **A)** Scatter plot of Log2 (fold change) in differential expression gene (DEG) analysis of P2 *Sox2+* cells comparing P2 *Sox2-* cells and E11.5 *Sox2+* cells comparing E11.5 *Sox2-* cells. **B)** Scatter plot of Log2(TPM+1) expression level of all transcription factor comparing P2 *Sox2+* cells with E11.5 *Sox2+* cells. Abbreviation: TPM, Transcript Per Million; DL, dental lamina; DESC, dental epithelial stem cell.

To systematically evaluate the dynamacis of transcription factors expression change between E11.5 *Sox2+* DL cells and P2 *Sox2+* DESC, we compared expression level of all transcription factors (**Fig. 2B**). We found there is strong corrlation between these two stages (majority of transcription factors have similar expression level in these two stages). In fact, if we divided all transcription factors into 4 groups based on their expression level measured by log2(TPM+1) value (TPM: transcripts per million), alluvial flow diagram shows that majority of them remain the same group in these two stages (**Fig. 3A**). Specifically, *Pitx2, Cnbp, Id3* and *Hmga1* were the highest expressed transcription factors in both stages (**Fig. 2B**). However, we found few transcription factors’ expression were dramatically reduced in P2 *Sox2+* DESC relative to E11.5 *Sox2+* DL, including *Lef1*, *Msx1* and *Meis2* (**Fig. 2B**). The reduction of *Lef1*, a critical transcription factor for Wnt/β-catenin signaling pathway, is particulary interesting because it has been shown that overexpression of *Lef1* increases DESC proliferation, expands labial cervical loop stem cell compartment and produces rapidly growing long tusk-like incisors in mice[38]. On the contrary, we found few transcription factors’ expression dramatically increase in P2 *Sox2+* DESC relative to E11.5 *Sox2+* DL, including *Nfix, Tbx1, Etv1, Zbtb20, Sox9, Sox10*, etc (**Fig. 2B**). We have previously found *Tbx1* is critical for ameloblast cell lineage differentiation[39, 40].

**Figure 3.**
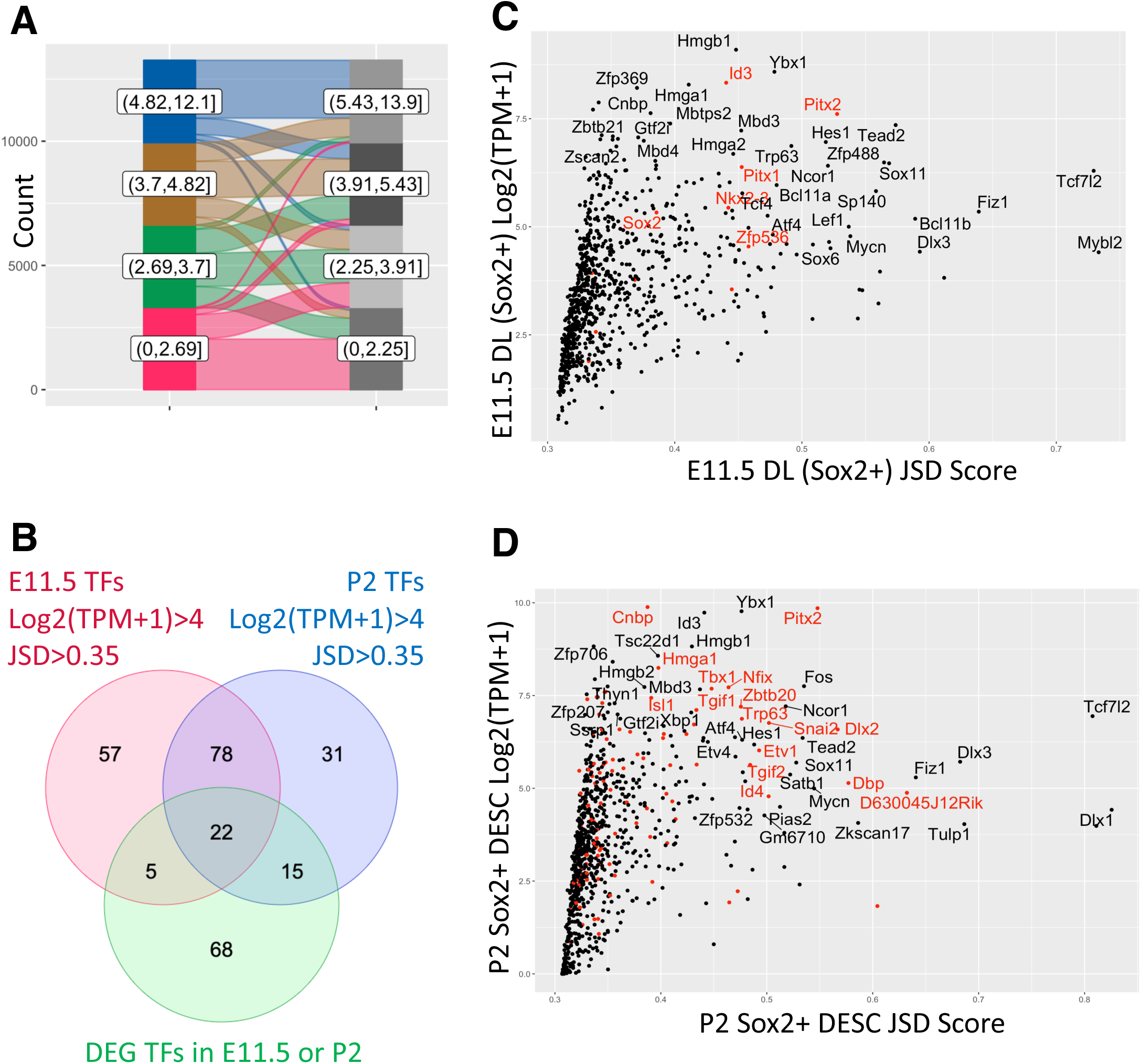
Systematically identify potential key transcription factors for P2 *Sox2+* DESC and E11.5 *Sox2+* DL cells. **A)** Alluvial flow diagram shows the dynamic of all genes’ expression level as measured with Log2(TPM+1) between P2 Sox2+ DESC and E11.5 Sox2+ DL cells. **B)** Overlapping of transcription factors that filter for 3 different criteria for potential key transcription factors. **C)** Scatter plot of all transcription factors’ expression level and tissue specificity score in E11.5 DL *Sox2+* cells. **D)** Scatter plot of all transcription factors’ expression level and tissue specificity score in P2 Sox2+ DESC cells. that filter for 3 different criteria for potential key transcription factors. Abbreviation: TPM, Transcript Per Million; DL, dental lamina; DESC, dental epithelial stem cell. JSD: tissue specificity score based on Jensen-Shannon divergence.

It has been shown that a small number of so-called key/master transcription factors is sufficient to establish the gene expression programs that define cell identity. Futhermore, it has been shown that over-expression of few key/master transcription factors is sufficient to direct cell lineage conversion/reprogram[41]. It has been shown that key/master transcription factors not only have relatively high expression levels, but also are typically expressed in a cell-type-specific fashion[42]. A quantative score based on Jensen-Shannon divergence (JSD score) has been developed to quantifies the tissue specificity of transcription factor in the cell type of interest (relative to other cell types in reference background dataset) [42]. Higer JSD score (closer to 1.0) means this transcription factor is more specific/enriched in the cell type of interest. To identify potential key/master transcription factors for dental epithelial stem cell, we calculated JSD score for both E11.5 *Sox2+* DL cells and P2 *Sox2+* DESC with their transcriptome data. We generated scatter plot with JSD score and Log2(TPM+1) expression value to identify transcription factors that not only have relatively high expression levels, but also are relative specific for dental epithelial stem cell (**Fig. 3C, 3D**). We found in both stages, majority of transcription factors have JSD score less than 0.35; only small fraction of transcription factors have JSD score higher than 0.35 (**Fig. 3C, 3D**). Specifically, in E11.5 *Sox2+* DL cells, we found 162 transcription factors that have JSD score higher than 0.35 and log2(TPM+1) expression value above 4.0 (around the median of all gene expression value, **Fig. 3A**) (**Fig. 3C**). Similarly, we found 146 transcription factors that have JSD score higher than 0.35 and log2(TPM+1) expression value above 4.0 in P2 *Sox2+* DESC (**Fig. 3D**). Furthermore, we found 100 transcription factors were shared in these two sets (**Fig. 3B**). We further filtered these 100 transcription factors based on DEG analysis results comparing *Sox2+* cells with *Sox2-* cells at E11.5 & P2. We identified potential 22 key/master transcription factors that meet following criteria: 1) highly expressed in both E11.5 *Sox2+* DL cells and P2 *Sox2+* DESC (Log2(TPM+1) expression value above 4.0); 2) highly specific for E11.5 *Sox2+* DL cells and P2 *Sox2+* DESC (JSD score above 0.35); and 3) significantly upregulated in at leaset one stage DEG analysis comparing *Sox2+* cells with *Sox2-* cells (**Fig. 3B**).

It has recently been proposed that cell identify and cell type specific transcription regulatory networks are defined by a group of special enhancers, so-called super-enhancers (SEs), are large clusters of enhancers with aberrant high levels of transcription factor binding[43–45]. Furthermore, it has been shown that many key/master transcription factors’ expression are driven by super-enhancers and binding motifs of many key/master transcription factors are found enriched in these super-enhancers. Super-enhancers are typically identified by analyzing H3K27ac ChIP-seq data with the ROSE algorithm. However, recently developed ATAC-Seq can be used to identify active enhancers and super-enhancers instead of H3K27ac ChIP-seq[46, 47]. ATAC-Seq protocol need much less cells comparing with traditional ChIP-seq (ATAC-seq has been successfully applied to as few as 500 cells[34, 48]). We performed ATAC-seq for ectoderm cell of E11.5 first pharyngeal arch (PA1, including maxillary process and a mandibular process). Additionally, we performed ATAC-seq for P2 *Sox2+* DESC isolated from labial cervical loop (same procedure as RNA-seq shown in **Fig. 1A**). We then identified super-enhancers with ROSE algorithm. We identified 2405 super-enhancers in E11.5 PA1 ectoderm (**Fig. 4A**), and 1366 super-enhancers in P2 *Sox2+* DESC (**Fig. 4B**). Importantly, we found that many potential key transcription factors identified with expression level and tissue specifity (**Fig. 3B**) were driven by super-enhancers including: *Pitx2, Id3, Pitx1, Tbx1, Trp63, Nkx2-3, Grhl3, Dlx2, Runx1, Nfix, Zfp536*, etc (**Fig. 4A, 4B**). Addtionally, motif enrichment analysis with HOMER found these transcription factors’ motif were significantly enriched in super-enhancers. These results further support that these few transcription factors might function as key transcription factors that define cell identity of *Sox2+* dental epithelial stem cells.

**Figure 4.**
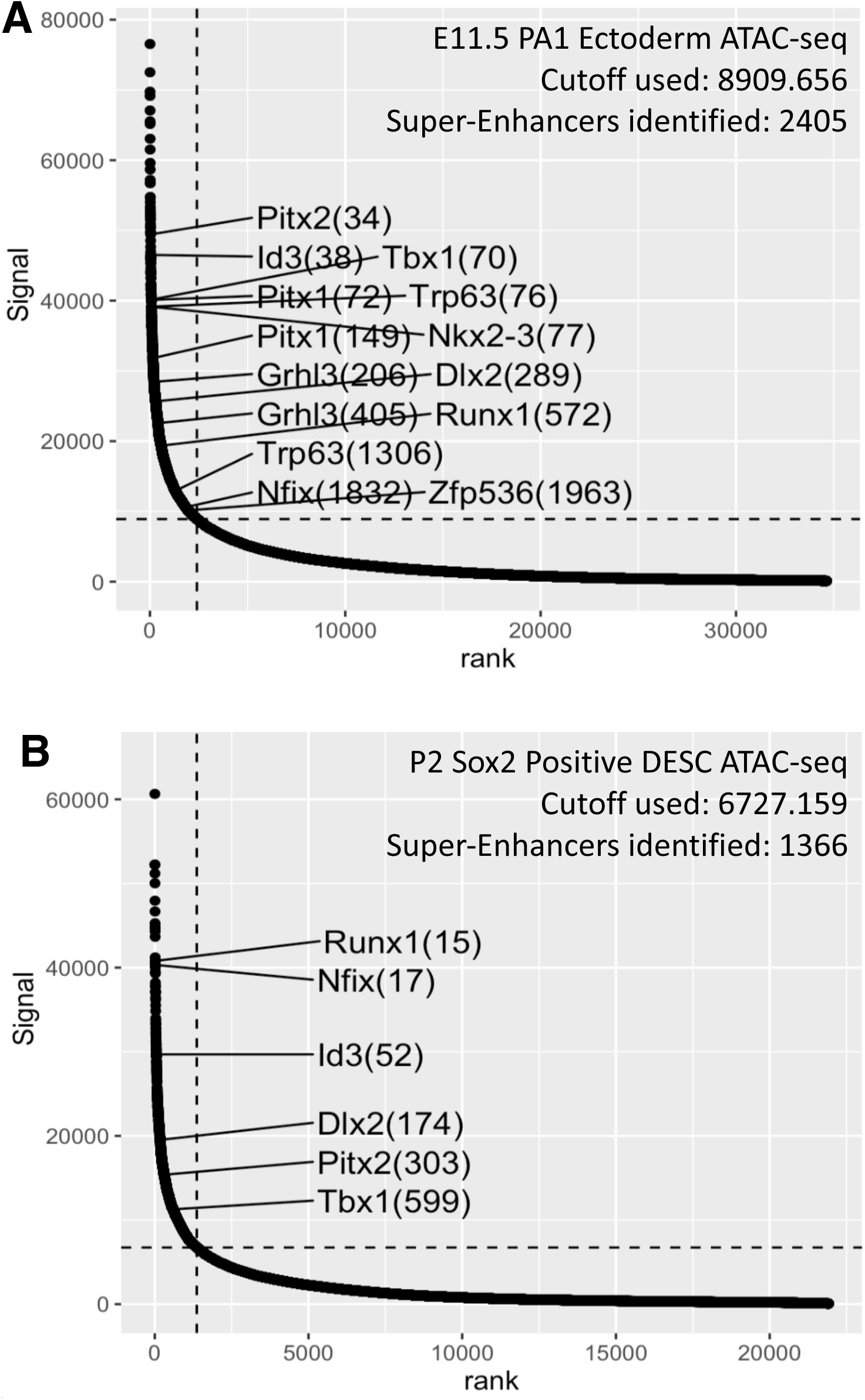
Key transcription factors for P2 *Sox2+* DESC and E11.5 *Sox2+* DL cells are driven super-enhancers. **A)** Identification of super-enhancers for E11.5 first pharyngeal arch (PA1, including maxillary process and a mandibular process). Selected super-enhancers with their assigned genes were marked with its ranking. **B)** Identification of super-enhancers for P2 Sox2+ DESC. Selected super-enhancers with their assigned genes were marked with its ranking.

## MATERIALS AND METHODS

### Mouse procedures

Mouse maintenance and mouse-related procedures were performed following protocols approved by the Institutional Animal Care and Use Committee of the University of Iowa. Embryos were staged by checking for vaginal plugs in the crossed females, with noon on the day a copulatory plug was present denoted as E0.5.

### Tissue preparation for RNA-seq and ATAC-seq

For ATAC-seq experiment, E11.5 mouse first pharyngeal arch epithelium were separated from mesenchyme with brief Dispase II (2mg/ml in PBS) treatment (20 mins @ 37°C) as previous described [49]. ATAC-seq libraries was prepared with the improved ATAC-seq protocol as described[34]. To isolate P2 Sox2+ dental epithelial stem cell, labial cervical loop region of P2 lower incisors were dissected. Single-Cell were prepared with cold protease protocol as previous described[50, 51]. 0.4% trypan blue was used for quality check. Based on GFP expression, we collected Sox2 positive cells and Sox2 negative cells. For RNA-seq experiment, Smart-seq2 protocol[33] was followed for cDNA preparation, Tn5 Tagmentation, and libraries preparation. Quality of cDNA and sequencing libraries were determined using a Bioanalyzer system (Agilent) and real time PCR. The KAPA Library Quantification Kit (Roche) was used for quantification of ATAC-seq and RNA-seq libraries.

### RNA-seq and ATAC-seq data analysis

RNA-seq and ATAC-seq reads were quality checked using the FastQC tool (http://www.bioinformatics.babraham.ac.uk/projects/fastqc). Low-quality and adapter sequences were removed using Trimmomatic[52]. For RNA-seq data, expression of transcripts was quantified using Salmon[53], and estimates of transcript abundance for gene-level analysis were imported and summarized using the tximport[54] function of the R/Bioconductor software suite[55]. Differentially expressed genes (DEGs) were identified by applying the R/Bioconductor package DeSeq2[35]. To calculate JSD tissue specifity score (based on Jensen-Shannon divergence), we used python package MTFinder (https://github.com/bioinfocao/MTFinder). For ATAC-seq data, reads were aligned to mm10 genome with bowtie2[56] using default setting with following parameters (--very-sensitive). For peak calling, we used Genrich (https://github.com/jsh58/Genrich) with following parameters (j -r -e chrM). To identify Super-enhancers based on ATAC-seq data, we used ROSE package with default setting (https://bitbucket.org/young_computation/rose). Motif enrichment analysis was performed with HOMER package[57] using the “findMotifsGenome.pl” program with default settings.

## DATA AVAILABILITY STATEMENT

RNA-seq and ATAC-seq data have been deposited to the Gene Expression Omnibus (GEO) under accession number GSE250073 and GSE250074.

## ACKNOWLEDGEMENTS

We are grateful to the Iowa Institute of Human Genetics (IIHG) Genomics Division (University of Iowa) for helping us with genomic sequencing. We also thank University of Iowa Research Services for providing High Performance Computing Resources. Additionally, we would like to thank members of the Cao and Van Otterloo laboratories for helpful discussion and suggestions. This study was supported by grants from National Institute of Dental and Craniofacial Research (R21DE029828 to H.C.; R01DE033009 to H.C. and E.V.O.) and seed grants from University of Iowa College of Dentistry and Dental Clinics (to H.C.).

## AUTHOR CONTRIBUTIONS

F.S.: performed experiments, data curation, data analysis, data interpretation, result visualization, prepare manuscript draft, manuscript review and editing; E.V.O.: data curation, data analysis, data interpretation, manuscript review and editing, and funding acquisition; H.C.: project conceptualization, project supervision, performed experiments, data curation, data analysis, data interpretation, result visualization, prepare manuscript draft, manuscript review and editing, and funding acquisition.

## Notes

### Competing Interest Statement

The authors have declared no competing interest.

## REFERENCE

1. Wu P, Wu X, Jiang T-X, Elsey RM, Temple BL, Divers SJ, et al. Specialized stem cell niche enables repetitive renewal of alligator teeth. PNAS. 2013;110(22):E2009–E18. doi: 10.1073/pnas.1213202110.

2. Handrigan GR, Richman JM. Autocrine and paracrine Shh signaling are necessary for tooth morphogenesis, but not tooth replacement in snakes and lizards (Squamata). Developmental Biology. 2010;337(1):171–86. doi: 10.1016/j.ydbio.2009.10.020.

3. Handrigan GR, Leung KJ, Richman JM. Identification of putative dental epithelial stem cells in a lizard with life-long tooth replacement. Development. 2010;137(21):3545–9. doi: 10.1242/dev.052415.

4. Huysseune A. Formation of a successional dental lamina in the zebrafish (Danio rerio): support for a local control of replacement tooth initiation. The International Journal of Developmental Biology. 2006;50(7):637–43. doi: 10.1387/ijdb.052098ah.

5. Smith MM, Fraser GJ, Mitsiadis TA. Dental lamina as source of odontogenic stem cells: evolutionary origins and developmental control of tooth generation in gnathostomes. Journal of Experimental Zoology Part B: Molecular and Developmental Evolution. 2009;312B(4):260-80. doi: 10.1002/jez.b.21272.

6. Smith Moya M, Fraser Gareth J, Chaplin N, Hobbs C, Graham A. Reiterative pattern of sonic hedgehog expression in the catshark dentition reveals a phylogenetic template for jawed vertebrates. Proceedings of the Royal Society B: Biological Sciences. 2009;276(1660):1225-33. doi: 10.1098/rspb.2008.1526.

7. Abduweli D, Baba O, Tabata MJ, Higuchi K, Mitani H, Takano Y. Tooth replacement and putative odontogenic stem cell niches in pharyngeal dentition of medaka (Oryzias latipes ). Microscopy. 2014;63(2):141–53. doi: 10.1093/jmicro/dft085.

8. Rasch LJ, Martin KJ, Cooper RL, Metscher BD, Underwood CJ, Fraser GJ. An ancient dental gene set governs development and continuous regeneration of teeth in sharks. Developmental Biology. 2016;415(2):347–70. doi: 10.1016/j.ydbio.2016.01.038.

9. Martin KJ, Rasch LJ, Cooper RL, Metscher BD, Johanson Z, Fraser GJ. Sox2+ progenitors in sharks link taste development with the evolution of regenerative teeth from denticles. PNAS. 2016;113(51):14769–74. doi: 10.1073/pnas.1612354113.

10. Handrigan GR, Richman JM. A network of Wnt, hedgehog and BMP signaling pathways regulates tooth replacement in snakes. Developmental Biology. 2010;348(1):130–41. doi: 10.1016/j.ydbio.2010.09.003.

11. Miura M, Gronthos S, Zhao M, Lu B, Fisher LW, Robey PG, Shi S. SHED: stem cells from human exfoliated deciduous teeth. Proc Natl Acad Sci U S A. 2003;100(10):5807–12. Epub 20030425. doi: 10.1073/pnas.0937635100. PubMed PMID: 12716973; PubMed Central PMCID: PMCPMC156282.

12. Morsczeck C, Gotz W, Schierholz J, Zeilhofer F, Kuhn U, Mohl C, et al. Isolation of precursor cells (PCs) from human dental follicle of wisdom teeth. Matrix Biol. 2005;24(2):155–65. Epub 20050212. doi: 10.1016/j.matbio.2004.12.004. PubMed PMID: 15890265.

13. Gronthos S, Mankani M, Brahim J, Robey PG, Shi S. Postnatal human dental pulp stem cells (DPSCs) in vitro and in vivo. Proc Natl Acad Sci U S A. 2000;97(25):13625–30. doi: 10.1073/pnas.240309797. PubMed PMID: 11087820; PubMed Central PMCID: PMCPMC17626.

14. Seo BM, Miura M, Gronthos S, Bartold PM, Batouli S, Brahim J, et al. Investigation of multipotent postnatal stem cells from human periodontal ligament. Lancet. 2004;364(9429):149-55. doi: 10.1016/S0140-6736(04)16627-0. PubMed PMID: 15246727.

15. Diaz A, Orozco J, Fonseca M. Multiple hyperodontia: report of a case with 17 supernumerary teeth with non syndromic association. Med Oral Patol Oral Cir Bucal. 2009;14(5):E229–31. Epub 20090501. PubMed PMID: 19218904.

16. Wang XP, Fan J. Molecular genetics of supernumerary tooth formation. Genesis. 2011;49(4):261–77. Epub 20110401. doi: 10.1002/dvg.20715. PubMed PMID: 21309064; PubMed Central PMCID: PMCPMC3188466.

17. Jernvall J, Thesleff I. Tooth shape formation and tooth renewal: evolving with the same signals. Development. 2012;139(19):3487–97. doi: 10.1242/dev.085084.

18. Shao F, Phan A-V, Van Otterloo E, Cao H. Transcriptional programs controlling lineages specification of mandibular epithelium during tooth initiation. bioRxiv. 2022:2022.11.26.518052. doi: 10.1101/2022.11.26.518052.

19. Juuri E, Saito K, Ahtiainen L, Seidel K, Tummers M, Hochedlinger K, et al. Sox2+ Stem Cells Contribute to All Epithelial Lineages of the Tooth via Sfrp5+ Progenitors. Developmental Cell. 2012;23(2):317–28. doi: 10.1016/j.devcel.2012.05.012.

20. Sanz-Navarro M, Seidel K, Sun Z, Bertonnier-Brouty L, Amendt BA, Klein OD, Michon F. Plasticity within the niche ensures the maintenance of a Sox2+ stem cell population in the mouse incisor. Development. 2018;145(1):dev155929. doi: 10.1242/dev.155929.

21. Seidel K, Ahn CP, Lyons D, Nee A, Ting K, Brownell I, et al. Hedgehog signaling regulates the generation of ameloblast progenitors in the continuously growing mouse incisor. Development. 2010;137(22):3753–61. doi: 10.1242/dev.056358. PubMed PMID: 20978073; PubMed Central PMCID: PMCPMC3049275.

22. Seidel K, Marangoni P, Tang C, Houshmand B, Du W, Maas RL, et al. Resolving stem and progenitor cells in the adult mouse incisor through gene co-expression analysis. Elife. 2017;6. Epub 20170505. doi: 10.7554/eLife.24712. PubMed PMID: 28475038; PubMed Central PMCID: PMCPMC5419740.

23. Biehs B, Hu JK, Strauli NB, Sangiorgi E, Jung H, Heber RP, et al. BMI1 represses Ink4a/Arf and Hox genes to regulate stem cells in the rodent incisor. Nat Cell Biol. 2013;15(7):846–52. Epub 20130602. doi: 10.1038/ncb2766. PubMed PMID: 23728424; PubMed Central PMCID: PMCPMC3735916.

24. Sharir A, Marangoni P, Zilionis R, Wan M, Wald T, Hu JK, et al. A large pool of actively cycling progenitors orchestrates self-renewal and injury repair of an ectodermal appendage. Nat Cell Biol. 2019;21(9):1102–12. Epub 20190902. doi: 10.1038/s41556-019-0378-2. PubMed PMID: 31481792; PubMed Central PMCID: PMCPMC6935352.

25. Juuri E, Jussila M, Seidel K, Holmes S, Wu P, Richman J, et al. Sox2 marks epithelial competence to generate teeth in mammals and reptiles. Development. 2013;140(7):1424–32. doi: 10.1242/dev.089599.

26. Kim EJ, Jung SY, Wu Z, Zhang S, Jung HS. Sox2 maintains epithelial cell proliferation in the successional dental lamina. Cell Prolif. 2020;53(1):e12729. Epub 20191119. doi: 10.1111/cpr.12729. PubMed PMID: 31746095; PubMed Central PMCID: PMCPMC6985665.

27. Square TA, Sundaram S, Mackey EJ, Miller CT. Distinct tooth regeneration systems deploy a conserved battery of genes. Evodevo. 2021;12(1):4. Epub 20210325. doi: 10.1186/s13227-021-00172-3. PubMed PMID: 33766133; PubMed Central PMCID: PMCPMC7995769.

28. Grieco TM, Richman JM. Coordination of bilateral tooth replacement in the juvenile gecko is continuous with in ovo patterning. Evol Dev. 2018;20(2):51–64. Epub 20180110. doi: 10.1111/ede.12247. PubMed PMID: 29318754; PubMed Central PMCID: PMCPMC5834371.

29. Chiba Y, Saito K, Martin D, Boger ET, Rhodes C, Yoshizaki K, et al. Single-Cell RNA-Sequencing From Mouse Incisor Reveals Dental Epithelial Cell-Type Specific Genes. Front Cell Dev Biol. 2020;8:841. Epub 20200901. doi: 10.3389/fcell.2020.00841. PubMed PMID: 32984333; PubMed Central PMCID: PMCPMC7490294.

30. Hermans F, Bueds C, Hemeryck L, Lambrichts I, Bronckaers A, Vankelecom H. Establishment of inclusive single-cell transcriptome atlases from mouse and human tooth as powerful resource for dental research. Front Cell Dev Biol. 2022;10:1021459. Epub 20221010. doi: 10.3389/fcell.2022.1021459. PubMed PMID: 36299483; PubMed Central PMCID: PMCPMC9590651.

31. Fresia R, Marangoni P, Burstyn-Cohen T, Sharir A. From Bite to Byte: Dental Structures Resolved at a Single-Cell Resolution. J Dent Res. 2021;100(9):897–905. Epub 20210325. doi: 10.1177/00220345211001848. PubMed PMID: 33764175; PubMed Central PMCID: PMCPMC8293759.

32. Arnold K, Sarkar A, Yram MA, Polo JM, Bronson R, Sengupta S, et al. Sox2(+) adult stem and progenitor cells are important for tissue regeneration and survival of mice. Cell Stem Cell. 2011;9(4):317–29. Epub 2011/10/11. doi: 10.1016/j.stem.2011.09.001. PubMed PMID: 21982232; PubMed Central PMCID: PMCPMC3538360.

33. Picelli S, Bjorklund AK, Faridani OR, Sagasser S, Winberg G, Sandberg R. Smart-seq2 for sensitive full-length transcriptome profiling in single cells. Nat Methods. 2013;10(11):1096–8. Epub 2013/09/24. doi: 10.1038/nmeth.2639. PubMed PMID: 24056875.

34. Corces MR, Trevino AE, Hamilton EG, Greenside PG, Sinnott-Armstrong NA, Vesuna S, et al. An improved ATAC-seq protocol reduces background and enables interrogation of frozen tissues. Nat Methods. 2017;14(10):959–62. Epub 20170828. doi: 10.1038/nmeth.4396. PubMed PMID: 28846090; PubMed Central PMCID: PMCPMC5623106.

35. Love MI, Huber W, Anders S. Moderated estimation of fold change and dispersion for RNA-seq data with DESeq2. Genome Biol. 2014;15(12):550. Epub 2014/12/18. doi: 10.1186/s13059-014-0550-8. PubMed PMID: 25516281; PubMed Central PMCID: PMCPMC4302049.

36. Stephens M. False discovery rates: a new deal. Biostatistics. 2017;18(2):275–94. doi: 10.1093/biostatistics/kxw041. PubMed PMID: 27756721; PubMed Central PMCID: PMCPMC5379932.

37. Shen WK, Chen SY, Gan ZQ, Zhang YZ, Yue T, Chen MM, et al. AnimalTFDB 4.0: a comprehensive animal transcription factor database updated with variation and expression annotations. Nucleic Acids Res. 2022. Epub 20221021. doi: 10.1093/nar/gkac907. PubMed PMID: 36268869.

38. Sun Z, Yu W, Sanz Navarro M, Sweat M, Eliason S, Sharp T, et al. Sox2 and Lef-1 interact with Pitx2 to regulate incisor development and stem cell renewal. Development. 2016;143(22):4115–26. doi: 10.1242/dev.138883.

39. Cao H, Florez S, Amen M, Huynh T, Skobe Z, Baldini A, Amendt BA. Tbx1 regulates progenitor cell proliferation in the dental epithelium by modulating Pitx2 activation of p21. Dev Biol. 2010;347(2):289–300. Epub 2010/09/08. doi: 10.1016/j.ydbio.2010.08.031. PubMed PMID: 20816801; PubMed Central PMCID: PMCPMC3334818.

40. Caton J, Luder HU, Zoupa M, Bradman M, Bluteau G, Tucker AS, et al. Enamel-free teeth: Tbx1 deletion affects amelogenesis in rodent incisors. Dev Biol. 2009;328(2):493–505. Epub 20090220. doi: 10.1016/j.ydbio.2009.02.014. PubMed PMID: 19233155; PubMed Central PMCID: PMCPMC2880856.

41. Takahashi K, Yamanaka S. A decade of transcription factor-mediated reprogramming to pluripotency. Nat Rev Mol Cell Biol. 2016;17(3):183–93. Epub 20160217. doi: 10.1038/nrm.2016.8. PubMed PMID: 26883003.

42. D’Alessio AC, Fan ZP, Wert KJ, Baranov P, Cohen MA, Saini JS, et al. A Systematic Approach to Identify Candidate Transcription Factors that Control Cell Identity. Stem Cell Reports. 2015;5(5):763–75. doi: 10.1016/j.stemcr.2015.09.016. PubMed PMID: 26603904; PubMed Central PMCID: PMCPMC4649293.

43. Hnisz D, Abraham BJ, Lee TI, Lau A, Saint-Andre V, Sigova AA, et al. Super-enhancers in the control of cell identity and disease. Cell. 2013;155(4):934–47. Epub 20131010. doi: 10.1016/j.cell.2013.09.053. PubMed PMID: 24119843; PubMed Central PMCID: PMCPMC3841062.

44. Whyte WA, Orlando DA, Hnisz D, Abraham BJ, Lin CY, Kagey MH, et al. Master transcription factors and mediator establish super-enhancers at key cell identity genes. Cell. 2013;153(2):307–19. doi: 10.1016/j.cell.2013.03.035. PubMed PMID: 23582322; PubMed Central PMCID: PMCPMC3653129.

45. Adam RC, Yang H, Rockowitz S, Larsen SB, Nikolova M, Oristian DS, et al. Pioneer factors govern super-enhancer dynamics in stem cell plasticity and lineage choice. Nature. 2015;521(7552):366–70. Epub 20150318. doi: 10.1038/nature14289. PubMed PMID: 25799994; PubMed Central PMCID: PMCPMC4482136.

46. Adam RC, Yang H, Ge Y, Lien WH, Wang P, Zhao Y, et al. Temporal Layering of Signaling Effectors Drives Chromatin Remodeling during Hair Follicle Stem Cell Lineage Progression. Cell Stem Cell. 2018;22(3):398–413 e7. Epub 20180111. doi: 10.1016/j.stem.2017.12.004. PubMed PMID: 29337183; PubMed Central PMCID: PMCPMC6425486.

47. Perez-Rico YA, Boeva V, Mallory AC, Bitetti A, Majello S, Barillot E, Shkumatava A. Comparative analyses of super-enhancers reveal conserved elements in vertebrate genomes. Genome Res. 2017;27(2):259–68. Epub 20161213. doi: 10.1101/gr.203679.115. PubMed PMID: 27965291; PubMed Central PMCID: PMCPMC5287231.

48. Quillien A, Abdalla M, Yu J, Ou J, Zhu LJ, Lawson ND. Robust Identification of Developmentally Active Endothelial Enhancers in Zebrafish Using FANS-Assisted ATAC-Seq. Cell Rep. 2017;20(3):709–20. doi: 10.1016/j.celrep.2017.06.070. PubMed PMID: 28723572; PubMed Central PMCID: PMCPMC5562042.

49. Van Otterloo E, Milanda I, Pike H, Thompson JA, Li H, Jones KL, Williams T. AP-2alpha and AP-2beta cooperatively function in the craniofacial surface ectoderm to regulate chromatin and gene expression dynamics during facial development. Elife. 2022;11. Epub 2022/03/26. doi: 10.7554/eLife.70511. PubMed PMID: 35333176; PubMed Central PMCID: PMCPMC9038197.

50. Sekiguchi R, Hauser B. Preparation of Cells from Embryonic Organs for Single-Cell RNA Sequencing. Curr Protoc Cell Biol. 2019;83(1):e86. Epub 20190408. doi: 10.1002/cpcb.86. PubMed PMID: 30957983; PubMed Central PMCID: PMCPMC6506382.

51. Denisenko E, Guo BB, Jones M, Hou R, de Kock L, Lassmann T, et al. Systematic assessment of tissue dissociation and storage biases in single-cell and single-nucleus RNA-seq workflows. Genome Biol. 2020;21(1):130. Epub 20200602. doi: 10.1186/s13059-020-02048-6. PubMed PMID: 32487174; PubMed Central PMCID: PMCPMC7265231.

52. Bolger AM, Lohse M, Usadel B. Trimmomatic: a flexible trimmer for Illumina sequence data. Bioinformatics. 2014;30(15):2114–20. Epub 20140401. doi: 10.1093/bioinformatics/btu170. PubMed PMID: 24695404; PubMed Central PMCID: PMCPMC4103590.

53. Patro R, Duggal G, Love MI, Irizarry RA, Kingsford C. Salmon provides fast and bias-aware quantification of transcript expression. Nat Methods. 2017;14(4):417–9. Epub 20170306. doi: 10.1038/nmeth.4197. PubMed PMID: 28263959; PubMed Central PMCID: PMCPMC5600148.

54. Soneson C, Love MI, Robinson MD. Differential analyses for RNA-seq: transcript-level estimates improve gene-level inferences. F1000Res. 2015;4:1521. Epub 20151230. doi: 10.12688/f1000research.7563.2. PubMed PMID: 26925227; PubMed Central PMCID: PMCPMC4712774.

55. Huber W, Carey VJ, Gentleman R, Anders S, Carlson M, Carvalho BS, et al. Orchestrating high-throughput genomic analysis with Bioconductor. Nat Methods. 2015;12(2):115–21. doi: 10.1038/nmeth.3252. PubMed PMID: 25633503; PubMed Central PMCID: PMCPMC4509590.

56. Langmead B, Trapnell C, Pop M, Salzberg SL. Ultrafast and memory-efficient alignment of short DNA sequences to the human genome. Genome Biol. 2009;10(3):R25. Epub 20090304. doi: 10.1186/gb-2009-10-3-r25. PubMed PMID: 19261174; PubMed Central PMCID: PMCPMC2690996.

57. Heinz S, Benner C, Spann N, Bertolino E, Lin YC, Laslo P, et al. Simple combinations of lineage-determining transcription factors prime cis-regulatory elements required for macrophage and B cell identities. Mol Cell. 2010;38(4):576–89. doi: 10.1016/j.molcel.2010.05.004. PubMed PMID: 20513432; PubMed Central PMCID: PMCPMC2898526.

